# Differential Expression Analysis of ZIKV Infected Human RNA Sequence Reveals Potential Biomarkers

**DOI:** 10.1101/498295

**Authors:** Almas Jabeen, Nadeem Ahmad, Khalid Raza

## Abstract

Zika virus (ZIKV) is considered to be an emerging viral outbreak due to its link to diseases like microcephaly, Guillain-Barre Syndrome in human. In this paper, we identify differentially expressed genes (DEGs) using RNA-seq data. In this study, we adopted the RNA-seq analysis pipeline to quantify RNA-seq data into read counts. Our analysis uncovers the significant DEGs which may be involved in the altered biological process somehow. Here, we report the list of significant DEGs, out of which three genes are found to be highly differentially expressed. In addition, our analysis also predicts other moderate DEGs, low DEGs whose differential expression was induced due to ZIKV infections.

## Introduction

According to the recently published reports by WHO, several types of diseases and infections outbreak all around the world of which majority were concerned with viral infections. Few of these are Ebola Virus Disease (EVD) [1], Middle East Respiratory Syndrome (MERS), H5N1 influenza infection and ZIKV infection [2]. During 2013 - 2014, the area under French Polynesia encountered the largest viral outbreak due to Zika [3] which grew to be of alarming concern as was a threat to the world. ZIKV is a flavivirus transmitted by Aedes mosquitoes [4]. It can be diagnosed via some of the molecular or other serologic testing involving blood, saliva (for neonates) [5], urine [6], etc. The minor infection results in lowgrade fever, myalgia, maculopapular rashes, and in severe cases, adults encounter neurological and congenital structural defects [7]. It may also cause congenital malformations in pregnant woman with fetal and newborn with microcephaly [8]. It was also found that Sertoli cells of mouse testis were susceptible to ZIKV infection, which implied that ZIKV can be transmitted sexually affecting human male reproductive system [9]. ZIKV also affects the eyes of adults mildly, but significantly in infants with infected mothers [7].

In [10], Xia et al showed that ZIKV fixes the mutations in NS1 gene that enhances mosquito infection and increases its ability to dodge immune response. The ZIKV infection also activates the P53 gene, causing genotoxic stress in the neural progenitors cells (NPCs) of human, similar to mutations as observed in genetically caused microcephaly and p53 [11]. ZIKV primarily targets the CD14 human blood monocytes inducing M2-skewed immune-suppression during pregnancy [12]. Autophagy process plays dual roles in ZIKV infection as both good and bad based on the viral replication stage [13]. ZIKV infection also affects peripheral nervous systems (PNS) as well as central nervous system, causing transcriptional dysregulation which results to cell death [14]. ZIKV depletes the NPCs in human cerebral organoids by triggering TLR3 receptor for innate immunity [15]. Though there are some licensed vaccines for other flaviviruses, still emerging threat of ZIKV outbreak laid urgent need of developing preventive vaccines and treatments for infected patients. Several candidate vaccines have been developed and identified potent for this usage, but they are not ready for widespread use. Makhluf & Shresta [16] described the development of ZIKV vaccines by utilizing the past knowledge from other dengue and flavivirus involving viral vectors, DNA-RNA dependent vaccines, live attenuated virus, immunogenic viral epitopes predicted *in silico* and also inactivated virus. Mesci et al.[17] showed that an FDA approved compound called Sofosbuvir (SOF), effective for Hepatitis C (HCV), also prevented human NPCs from ZIKV infection-mediated cell death. It also prevented vertical transmission of infection from infected mother to fetus [17]. The analysis of the structures of ZIKV NS3 and NS5 revealed some conserved features which may be used for structure-based design of the antiviral compounds against ZIKV infection [8].

The Next Generation Sequencing (NGS) technologies like RNA-seq (RNA sequencing) find their application in the diagnostic virology (discovery, characterization and detection of viruses), antiviral drug and vaccine development, analysis of host-virus interaction, study of viral spread, etc. Due to being cost effective and having an improved turnaround time, NGS methods along with other analytical and clinical tests validation methods, can serve as essential diagnostic for viral spread [18]. RNA-seq technology uses the NGS method that gives the snapshot of RNA present in some genome sample along with its quantification at a given moment of time. It is preferred over the Microarray technology for gene expression studies [2] [19]. The research in RNA-seq includes studying the altered pathway during infection or disease, gene expression changes (differential expression analysis), etc. RNA-seq along with other NGS platforms (SNP, CGH arrays) can be used in the detection of CNV behind gene expression and novel genes behind tumorogenesis [20]. RNA-seq is cheaper, highly sensitive and faster than traditional sequencing methods, detects more transcripts than microarray technology [21]. Though RNA-seq analysis is considered as the standard expression profiling methodology, still easy, open and standard pipelines for performing this task by non-expert research community with different background is a major challenge. The read quantification and differentially analyzing the expression data based on high throughput NGS data, such as RNA-seq are highly appreciated. Mostly, the research interest lies in the comparison of the transcription result under different conditions, therefore the RNA-seq analysis studies can be categorized into Differential Gene Expression (DGE) studies where the transcriptional measure of each gene is compared between conditions, Differential Transcript Expression (DTE) studies and Differential Transcript/exon Usage (DTU) studies [22] [40].

In this study, we performed DEGs analysis of ZIKV exposed patient using RNA-seq data from the NGS. The identified DEGs from this pipeline may have key roles in significant biological processes and functional pathways related to the disease; hence this would enable us to search for putative vaccines for viral infections like ZIKV.

## Methodology

In this pipeline the RNA-seq dataset of ZIKV infected human induced pluripotent stem cells (hiPSCs) from human were retrieved from NCBI. Then after read quality check by FastQC [23], the data was preprocessed using Trimmomatic [24]. Reads were then mapped to reference human genome (hg38)/annotated human genome using aligner named Bowtie2 [25], the resulting file underwent for quantification of expression values using HTSeq-count [26]. It simply counts the aligned reads overlapping genomic regions from the mapping step, providing us with expression value counts. These counts further serve as input for Normalization and DEGs identification. Bioconductor package of ‘R’ language [27] [28] provides several statistical tools such as edgeR [29] [30], DESeq2 [31] which performs Normalization. Additionally, Cuffdiff [32] [33] [34], an Ubuntu based program provided by Cufflinks tool, may also give DEGs. In this research, we identified the three DEGs using these three tools in a consensus manner.

The complete pipeline for our work is shown in Fig. 1, which includes various steps such as data collection, data preprocessing (quality check, adapter trimming, etc.), read mapping to the human genome, read counting, and differential expression analysis.

**Fig. 1.**
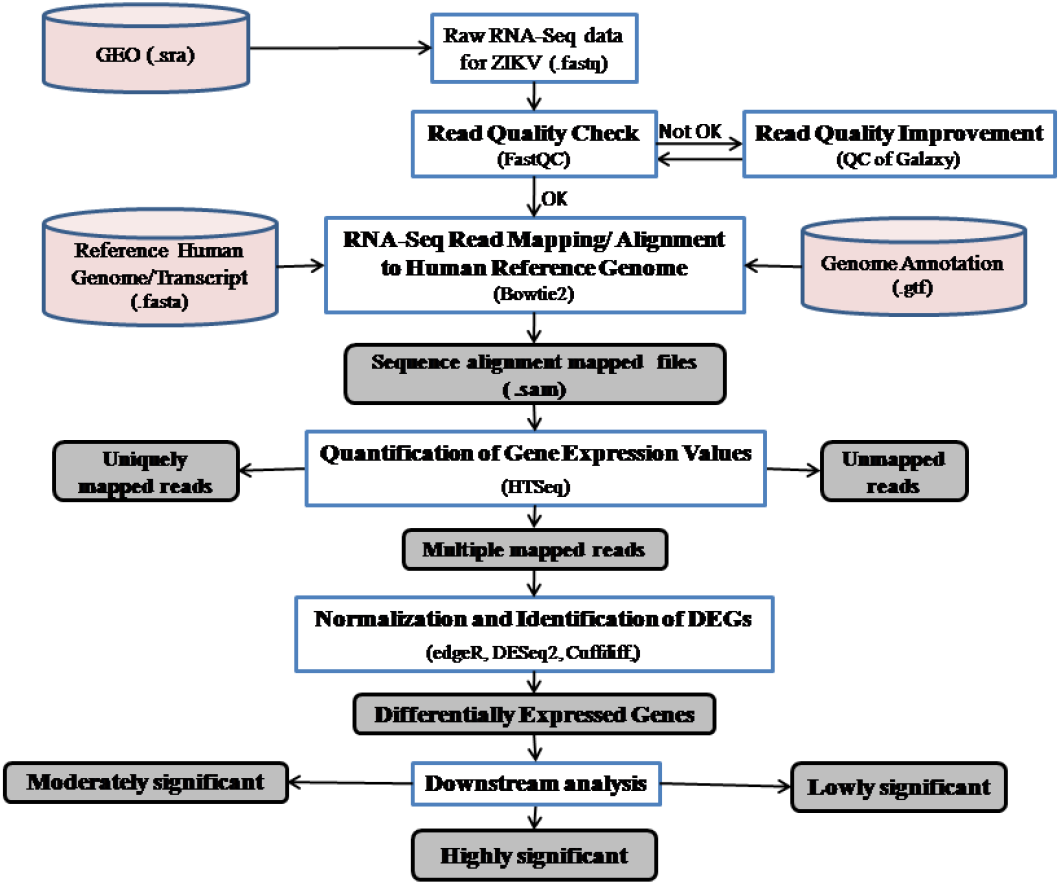
Workflowdiagram for Differentially Expressed Genes (DEGs) identification using ZIKV infection RNA-Seq data from the NGS.

### Raw data collection

GSE78711 GEO accession for ZikV infected hiPSCs-derived cortical neural progenitor cells (hNPCs) was retrieved from NCBI [35]. The data consist of eight sample files of which four were from control experiments (Mock) and other four from treated (ZIKV), i.e. virus treated experimentally.

### Data Preprocessing

#### Data format conversion (.sra to. fastq)

All the eight SRA files (.sra) were converted to FastQ format (.fastq) from ubuntu command prompt.

#### Raw data quality check and adaptor trimming

All the samples were quality checked using FastQC tool. We used Trimmomatic 0.36 tool for adapter trimming in order to remove the adapters from data.

#### Separation of all types of rRNAs

In order to segregate rRNAs from the sequence, we used SortMeRNA 2.1 tool [36].

### Read Mapping

We used Bowtie2 aligning tool for mapping the pre-processed reads against the human reference genome (hg38), which outputs aligned reads as .sam files.

### Read Counting

In order to count the number of reads which were mapped to the human reference genome, we used HTSeq-count tool. It takes aligned reads (.sam file) and a list of genome features (hg38) and provides read counts.

### Differential Expression Analysis

The differential expression analysis starts with the normalization step which is a method to adjust read counts between samples in such a way to get a uniform normalized expression values throughout the experiment. We applied following three tools.

#### edgeR

The edgeR [29] [30] is the expression analysis tool which models the mapped read count data using a negative binomial (NB) model. It moderates the estimated dispersion calculated for each gene to a single common dispersion estimate, or to a local dispersion estimate, which results from genes with similar expression weight calculated using a weighted conditional likelihood method [31]. It is a measure of assessing the inter-library variation of that gene.

For the classic edgeR analysis, we took eight sample libraries in two groups (group factors marked as 1 for control experiments and 2 for virus-treated experiments), and counts were stored in a tab-delimited text file with gene symbols in a column. After dispersion estimation, we performed exactTest for determining differential expression. On normalized expression values, we applied following three tools.

#### DESeq2

The DESeq2 package uses the NB model in order to test the differential expression. It estimates the shrinkage according to the data distribution and adjusts the logarithmic fold changes to improvise the result stability and interpretation [31]. For analysis through DESeq2 package, we input two files, one with count data of all eight samples in the form of a matrix of integer values, and other with the specified sample condition whether samples are controlled or virus-treated. It firstly estimates size factors, and then calculates gene-wise dispersion. It finally fits the model and tests for differential expression.

#### Cuffdiff

Cuffdiff [32] [33] [34] is Ubuntu based Cufflink transcript assembly package used to identify significant changes in transcript expression. It models the variance in groups of samples which lies beyond the expected variance calculated by Poisson model. It tests for observed logfold change in its expression against the null hypothesis of no change. The normalization process in Cuffdiff is performed by clas-sic-fpkm, geometric mode, quartile mode. Cuffdiff needs count files in .bam format, so firstly it was converted to. bam files using Samtools. After sorting it, ran Cuffdiff command.

### Consensus approach to DEGs

We assume that A is the set of DEGs identified by tool edgeR under the specified cutoffs and filters, B is the set of DEGs identified by tool DESeq2 and C is the set of DEGs identified by Cuffdiff tool. Using a consensus approach, differentially expressed highly significant genes (DEGsHigh) can be defined as Equation (1).

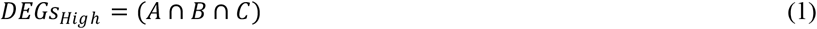

Similarly, using majority voting rule, differentially expressed moderately significant genes (DEGsModerate) can be defined as Equation (2),

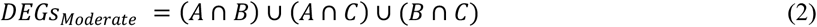

And differentially expressed low significant DEGs (DEGsLow) can be defined as Equation (3),

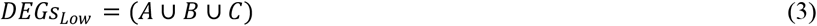

## Results and Discussion

The complete protocol as depicted in Figure 1 has been implemented and carried out on a workstation with 8 GB RAM, multi-core processors under Ubuntu 18.04.1 LTS operating system.

### Data Preprocessing

On read quality check of raw data by FastQC, we found that four out of eight samples had quality-score above 20 (Phred Score > 20), which were considered to be good whereas remaining four samples had it below 20. Poor quality reads when subjected for adaptor trimming; only 2-3% of the total reads got trimmed. All types of rRNAs were separated from the samples.

### Read Mapping

After read mapping to the human genome, a .sam file for each of the eight input files was obtained. We observed that the overall alignment rate of all the eight samples is more than 99%, which is considered to be very well aligned.

### Read counting and Normalization

The number of reads mapped (read count) to a gene is considered to be the proxy of its expression. For the data analysis, read count data from HTSeq-count further needs to be normalized by total fragment count in order to make counts comparable across the experiments. edgeR, DESeq2 and Cuffdiff were used for this task, which firstly transformed read count data into a continuous distribution. They used the NB model to estimate dispersion parameter for each gene. This dispersion parameter gives a measure of the degree of inter-library variation of that gene between the samples. Estimation of common dispersion provided the idea of overall variability across the genome dataset.

### Differential Expression Analysis

When all the eight samples grouped in two groups were subjected for differential analysis through the edgeR tool of R Bioconductor, it was found that normalization factors calculated for each sample is close to 1 which signifies that all the eight libraries are similar in composition. The input estimated common dispersion before estimating tagwise dispersions in order to proceed with differential expression analysis. Firstly, BCV was applied to this input data.

The Biological coefficient of variance (BCV) is the mathematical square root of common dispersion estimated using the NB model. It is the coefficient of variation in which the unknown true abundance of the gene varies between the samples. With an increase in the number of read counts, the BCV remains unaffected, though a decrease in technical CV is observed. Therefore, the accurate BCV estimation is crucial for differential expression analysis studies in RNA-seq experiments. The BCV calculated from the experiment was found to be 34%. Higher the BCV measure, lower will be the number of differentially expressed genes detected. In Fig. 2, a common dispersion (red line on BCV plot) lies between 0.2-0.4, hence considered to be detected higher number of DEGs.

**Fig. 2.**
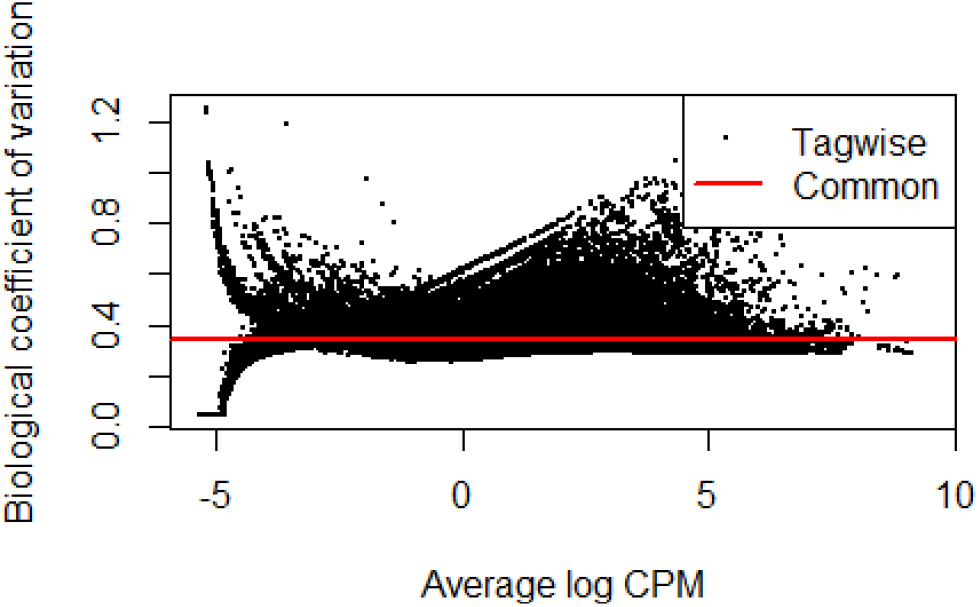
Plot for Biological coefficient of variation

For Multidimensional Scaling (MDS), the input was provided in the form of a distance matrix where values represent the distances between the pairs of objects. MDS plot represents the relationship between different groups of samples and can be affected by high BCVs. MDS plots show distances between the samples in terms of BCV as shown in Fig. 3.

**Fig. 3.**
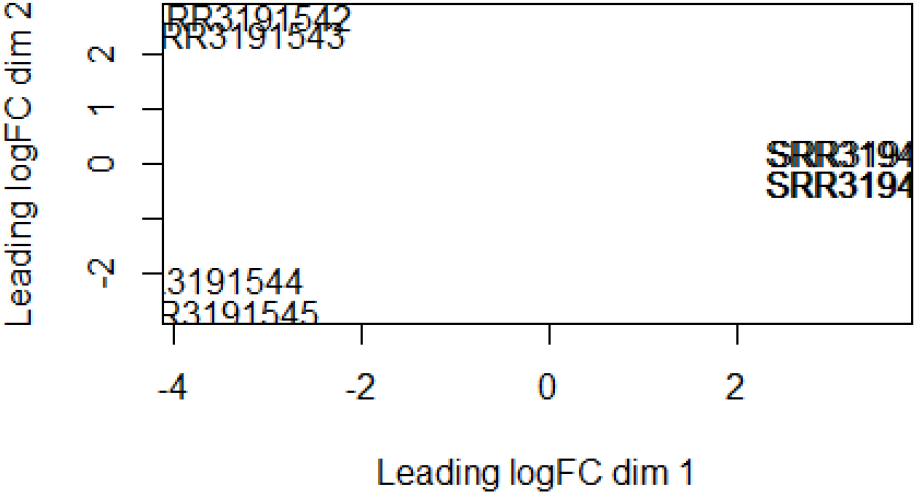
MDS plot of various samples.

Dimension 1 (dim1) separates the control samples from the Virus-treated samples which signify the possibility of detecting higher number of DEGs. This plot can be observed in the form of an unsupervised clustering.

After fitting the NB models and estimation of dispersions, we proceeded with tests determining the differential expression. The tagwise exactTest (similar to Fisher’s exact test) was applied. P values were calculated by combining over all sums of counts that have a probability less than the probability under the null hypothesis of the observed sum of counts. The test performed at FDR< 0.05, provided us with output result with all transcripts arranged in tabular form with information regarding their geneID, logFC, logCPM, P-value. It was observed that 683 genes were down-regulated, 152668 genes were not differentially expressed and 375 were up-regulated.

The smear plot of tagwise log-fold changes (logFC) against logCPM is shown in Fig. 4. The differentially expressed tags are highlighted in the plot and the horizontal blue lines show 4-fold changes.

**Fig. 4.**
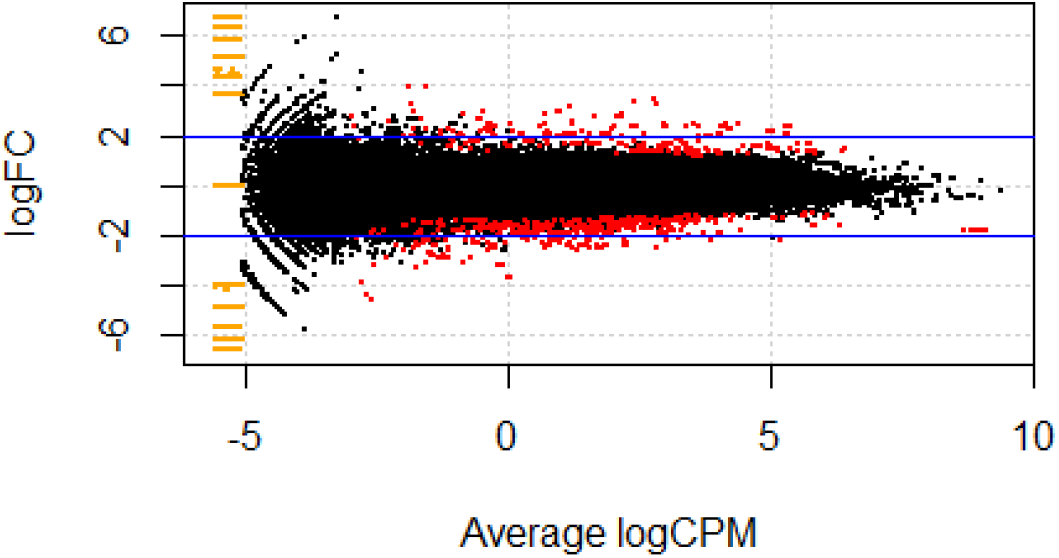
Smear plot of dataset analogous to an MA-plot as for microarray data

For the consensus method in DEGs identification, we also applied DESeq2 tool from R Bioconductor which gave the result along with normalized count data for all eight samples. The result constitutes the fields like log2 fold changes, p values, adjusted p values, etc. The dispersion plot of the normalized read count from DESeq2 tool is shown in Fig. 5(a).

**Fig. 5.**
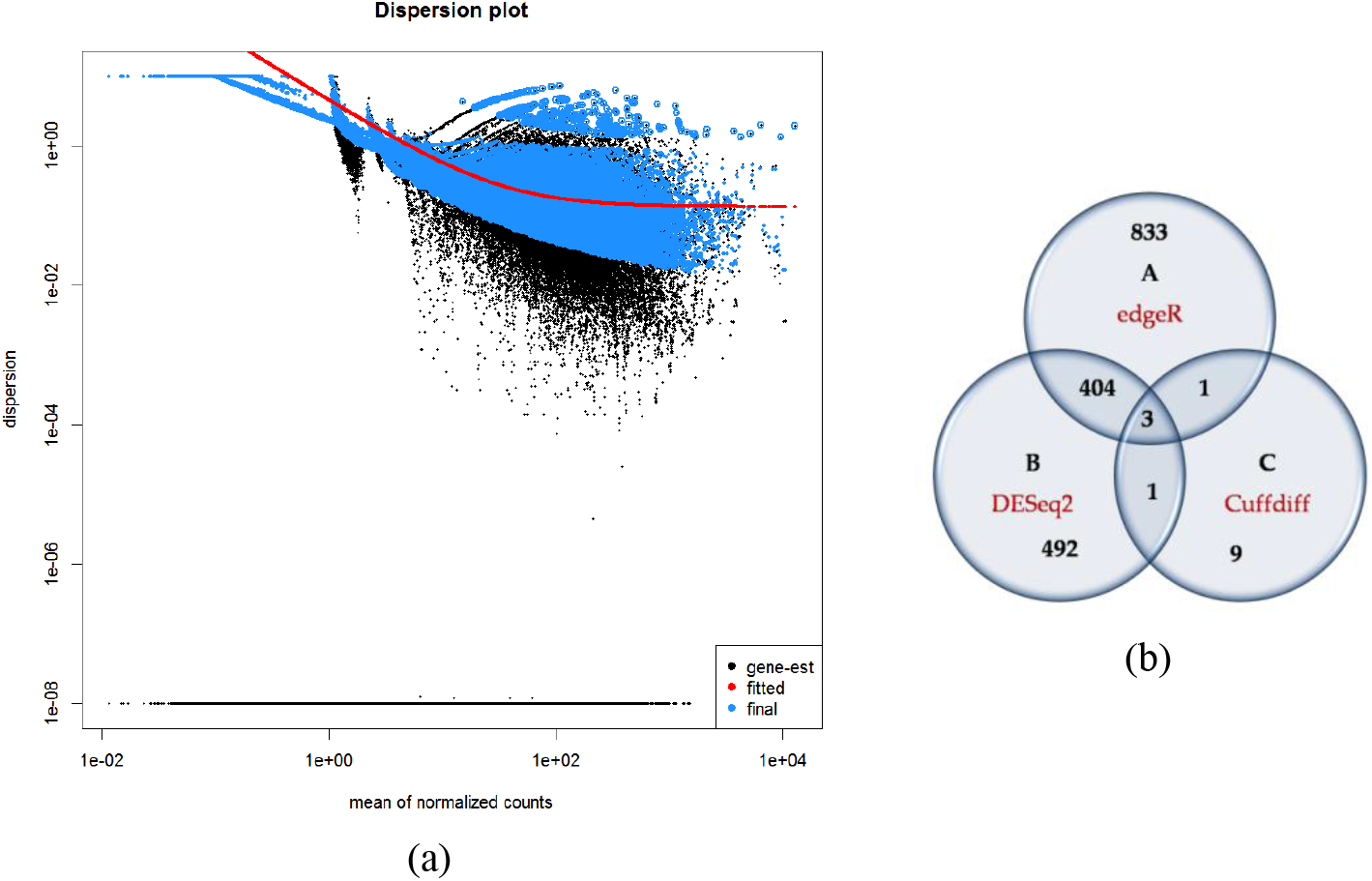
(a) Dispersion plot for mean of normalized read counts by DESeq2 tool. (b) Venn diagram representing number of identified DEGs from three tools.

We also performed a differential analysis of count data using Cuffdiff program. The processed input files from HTSeq tool when subjected to Cuffdiff tool, gave several output files for expression estimates in terms of FPKM and read counts. One of the files had identified DEGs in the form of a list.

Since results contain a large number of DEGs, we applied the filtering criteria to extract out best identified DEGs in an order from each tool. We classified the extracted significant DEGs into three categories, namely highly significant DEGs, moderately significant DEGs and low significant DEGs according to Equation (1), (2) and (3), respectively. In Fig.5(b) the number of DEGs identified by three different tools is shown.

Using a consensus approach, we found three DEGs were of high significance (NM_001010989.2, NM_001034.3, NM_001042.2), 406 were moderately significant and 513 were low significant (Fig. 6). The function of three identified highly significant DEGs are listed in Table 1. Out of these three genes, HERPUD1 have been reported in the literature to be involved somehow in ZIKV infections (PMID: 29915147, 28325921). However, list of moderately and low significant DEGs need further experimental validation for its reliability.

**Table 1.**
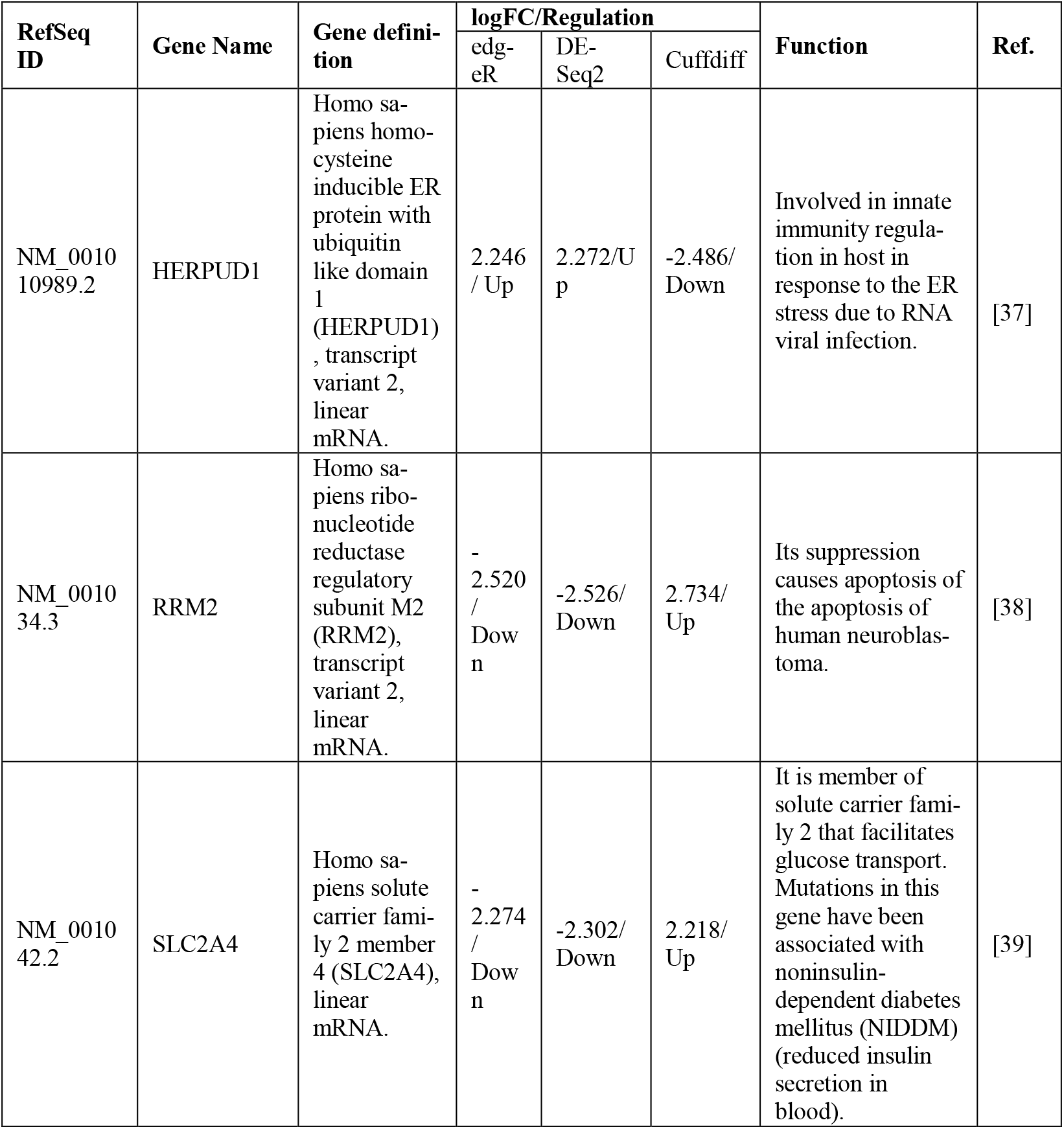
Functions of three identified highly significant genes.

| RefSeq ID | Gene Name | Gene definition | logFC/Regulation |  |  | Function | Ref. |
| --- | --- | --- | --- | --- | --- | --- | --- |
|  |  |  | edg-eR | DE-Seq2 | Cuffdiff |  |  |
| NM_001010989.2 | HERPUD1 | Homo sapiens homo-cysteine inducible ER protein with ubiquitin like domain 1 (HERPUD1), transcript variant 2, linear mRNA. | 2.246 / Up | 2.272/Up | -2.486/Down | Involved in innate immunity regulation in host in response to the ER stress due to RNA viral infection. | [37] |
| NM_001034.3 | RRM2 | Homo sapiens ribonucleotide reductase regulatory subunit M2 (RRM2), transcript variant 2, linear mRNA. | -2.520 / Down | -2.526/Down | 2.734/Up | Its suppression causes apoptosis of the apoptosis of human neuroblastoma. | [38] |
| NM_001042.2 | SLC2A4 | Homo sapiens solute carrier family 2 member 4 (SLC2A4), linear mRNA. | -2.274 / Down | -2.302/Down | 2.218/Up | It is member of solute carrier family 2 that facilitates glucose transport. Mutations in this gene have been associated with noninsulin-dependent diabetes mellitus (NIDDM) (reduced insulin secretion in blood). | [39] |

## Conclusion

The outbreak of viral disease such as ZIKV brought the attention of computational biologists and bioinformaticians to perform the differential expression analysis of ZIKV infected patients to understand transcriptomic changes in the body. Knowledge of transcriptomic changes helps the design of better diagnostic tools, therapeutics and treatments. In this study, we analyzed different expressions of ZIKV patients using RNA-seq data for NGS. We report three highly significant genes which are differentially expressed using a consensus of three tools (edgeR, DESeq2, Cuffdiff). These genes are validated from the literature for its reliability. In future work, we would perform pathway analysis, GO enrichment analysis, topological analysis of these three genes. We would also explore highly ranked genes which are moderately significant DEGs.

## Acknowledgement

The author Almas Jabeen acknowledges Maulana Azad National Fellowship-Junior Research Fellowship (JRF) received from the University Grant Commission, Government of India.

## References

1. Mudassar Imran, Adnan Khan, Ali R. Ansari, and Syed Touqeer Hussain Shah. Modeling transmission dynamics of Ebola virus disease. International Journal of Biomathematics, 10, no. 04: 1750057, 2017

2. Almas Jabeen, Nadeem Ahmad, and Khalid Raza. Machine Learning-Based State-of-the-Art Methods for the Classification of RNA-Seq Data. In Classification in BioApps, pp. 133–172 Springer, Cham, 2018. https://doi.org/10.1007/978-3-319-65981-7_6

3. Cao-Lormeau, Van-Mai, Alexandre Blake, Sandrine Mons, Stéphane Lastère, Claudine Roche, Jessica Vanhomwegen, Timothée Dub et al. Guillain-Barré Syndrome outbreak associated with Zika virus infection in French Polynesia: a case-control study. The Lancet, 387, no. 10027: 1531–1539, 2016.

4. Shashi K. Tiwari, Jason Dang, Yue Qin, Gianluigi Lichinchi, Vikas Bansal, and Tariq M. Rana. Zika virus infection reprograms global transcription of host cells to allow sustained infection.” Emerging microbes & infections, 6, no. 4: e24, 2017.

5. Didier Musso, Claudine Roche, Tu-Xuan Nhan, Emilie Robin, Anita Teissier, and Van-Mai Cao-Lormeau. Detection of Zika virus in saliva. Journal of Clinical Virology, 68: 53–55, 2015.

6. Ann-Claire Gourinat, Olivia O’Connor, Elodie Calvez, Cyrille Goarant, and Myrielle Dupont-Rouzeyrol. Detection of Zika virus in urine. Emerging infectious diseases, 21, no. 1: 84. 2015.

7. Rupesh Agrawal, Hnin Hnin Oo, Praveen K. Balne, Lisa Ng, Louis Tong, and Yee S. Leo. Zika virus and the eye. Ocular immunology and inflammation, 26, no. 5 (2018): 654–659. 2018.

8. Yi Shi, and George F. Gao. Structural biology of the Zika virus. Trends in biochemical sciences, 42, no. 6: 443–456, 2017.

9. Zi-Yang Sheng, Na Gao, Zhao-Yang Wang, Xiao-Yun Cui, De-Shan Zhou, Dong-Ying Fan, Hui Chen, Pei-Gang Wang, and Jing An. Sertoli cells are susceptible to ZIKV infection in mouse testis. Frontiers in cellular and infection microbiology, 7: 272, 2017.

10. Hongjie Xia, Huanle Luo, Chao Shan, Antonio E. Muruato, Bruno TD Nunes, Daniele BA Medeiros, Jing Zou et al. An evolutionary NS1 mutation enhances Zika virus evasion of host interferon induction. Nature communications, 9, no. 1: 414, 2018.

11. Vincent El Ghouzzi, Federico T. Bianchi, Ivan Molineris, Bryan C. Mounce, Gaia E. Berto, Malgorzata Rak, Sophie Lebon et al. ZIKA virus elicits P53 activation and genotoxic stress in human neural progenitors similar to mutations involved in severe forms of genetic microcephaly and p53. Cell death & disease, 8, no. 1: e2567, 2017.

12. Suan-Sin Foo, Weiqiang Chen, Yen Chan, James W. Bowman, Lin-Chun Chang, Younho Choi, Ji Seung Yoo et al. Asian Zika virus strains target CD14+ blood monocytes and induce M2-skewed immunosuppression during pregnancy. Nature microbiology, 2, no. 11: 1558, 2017.

13. Abhilash I. Chiramel and Sonja M. Best. Role of autophagy in Zika virus infection and pathogenesis. Virus research, 254: 34–40, 2018.

14. Yohan Oh, Feiran Zhang, Yaqing Wang, Emily M. Lee, In Young Choi, Hotae Lim, Fahimeh Mirakhori et al. Zika virus directly infects peripheral neurons and induces cell death. Nature neuroscience, 20, no. 9: 1209. 2017.

15. Jason Dang, Shashi K. Tiwari, Gianluigi Lichinchi, Yue Qin, Veena S. Patil, Alexey M. Eroshkin, and Tariq M. Rana. Zika virus depletes neural progenitors in human cerebral organoids through activation of the innate immune receptor TLR3. Cell stem cell 19, no. 2: 258–265. 2016.

16. Huda Makhluf and Sujan Shresta. Development of Zika virus vaccines. Vaccines 6, no. 1: 7,2018.

17. Pinar Mesci, Angela Macia, Spencer M. Moore, Sergey A. Shiryaev, Antonella Pinto, Chun-Teng Huang, Leon Tejwani et al. Blocking Zika virus vertical transmission. Scientific reports 8, no. 1: 1218. 2018.

18. Luisa Barzon, Enrico Lavezzo, Giulia Costanzi, Elisa Franchin, Stefano Toppo, and Giorgio Palù. Next-generation sequencing technologies in diagnostic virology. Journal of Clinical Virology, 58, no. 2: 346–350. 2013.

19. Ayman Grada and Kate Weinbrecht. Next-generation sequencing: methodology and application. The Journal of investigative dermatology 133, no. 8: e11, 2013.

20. Costa Valerio, Marianna Aprile, Roberta Esposito, and Alfredo Ciccodicola. RNA-Seq and human complex diseases: recent accomplishments and future perspectives. European Journal of Human Genetics 21, no. 2: 134. 2013.

21. Khalid Raza and Sabahuddin Ahmad. Recent advancement in next generation sequencing techniques and its computational analysis. International Journal of Bioinformatics Research and Applications (In press). [arXiv preprint arXiv:1606.05254]

22. Charlotte Soneson, Michael I. Love, and Mark D. Robinson. Differential analyses for RNA-seq: transcript-level estimates improve gene-level inferences. F1000Research 4, 2015.

23. Simon Andrews. FastQC: a quality control tool for high throughput sequence data, 2010.

24. Anthony M. Bolger, Marc Lohse and Bjoern Usadel. Trimmomatic: a flexible trimmer for Illumina sequence data. Bioinformatics 30, no. 15: 2114–2120, 2014.

25. Ben Langmead and Steven L. Salzberg. Fast gapped-read alignment with Bowtie 2. Nature methods 9, no. 4: 357, 2012.

26. Simon Anders, Paul Theodor Pyl, and Wolfgang Huber. HTSeq—a Python framework to work with high-throughput sequencing data. Bioinformatics 31, no. 2: 166–169. 2015.

27. Wolfgang Huber, Vincent J. Carey, Robert Gentleman, Simon Anders, Marc Carlson, Benilton S. Carvalho, Hector Corrada Bravo et al. Orchestrating high-throughput genomic analysis with Bioconductor. Nature methods 12, no. 2: 115. 2015.

28. Robert C. Gentleman, Vincent J. Carey, Douglas M. Bates, Ben Bolstad, Marcel Dettling, Sandrine Dudoit, Byron Ellis et al. Bioconductor: open software development for computational biology and bioinformatics. Genome biology 5, no. 10: R80, 2004.

29. Mark D. Robinson, Davis J. McCarthy and Gordon K. Smyth. edgeR: a Bioconductor package for differential expression analysis of digital gene expression data. Bioinformatics 26, no. 1: 139–140, 2010.

30. McCarthy, Davis J., Yunshun Chen, and Gordon K. Smyth. Differential expression analysis of multifactor RNA-Seq experiments with respect to biological variation. Nucleic acids research 40, no. 10: 4288–4297, 2012.

31. Michael I. Love, Wolfgang Huber and Simon Anders. Moderated estimation of fold change and dispersion for RNA-seq data with DESeq2. Genome biology 15, no. 12: 550, 2014.

32. Cole Trapnell, Brian A. Williams, Geo Pertea, Ali Mortazavi, Gordon Kwan, Marijke J. Van Baren, Steven L. Salzberg, Barbara J. Wold, and Lior Pachter. Transcript assembly and quantification by RNA-Seq reveals unannotated transcripts and isoform switching during cell differentiation. Nature biotechnology 28, no. 5: 511, 2010.

33. Cole Trapnell, David G. Hendrickson, Martin Sauvageau, Loyal Goff, John L. Rinn, and Lior Pachter. Differential analysis of gene regulation at transcript resolution with RNA-seq. Nature biotechnology 31, no. 1: 46, 2013.

34. Adam Roberts, Cole Trapnell, Julie Donaghey, John L. Rinn, and Lior Pachter. Improving RNA-Seq expression estimates by correcting for fragment bias. Genome biology 12, no. 3: R22, 2011.

35. Hengli Tang, Christy Hammack, Sarah C. Ogden, Zhexing Wen, Xuyu Qian, Yujing Li, Bing Yao et al. Zika virus infects human cortical neural progenitors and attenuates their growth. Cell stem cell 18, no. 5: 587–590, 2016.

36. Evguenia Kopylova, Laurent Noé and Héléne Touzet. SortMeRNA: fast and accurate filtering of ribosomal RNAs in metatranscriptomic data. Bioinformatics 28, no. 24: 3211–3217, 2012.

37. Maolin Ge, Zhen Luo, Zhi Qiao, Yao Zhou, Xin Cheng, Qibin Geng, Yanyan Cai et al. HERP Binds TBK1 To Activate Innate Immunity and Repress Virus Replication in Response to Endoplasmic Reticulum Stress. The Journal of Immunology: ji1700376, 2017.

38. Junfeng Li, Jinglin Pang, Yongdong Liu, Jing Zhang, Chuanguang Zhang, Gang Shen, and Lili Song. Suppression of RRM2 inhibits cell proliferation, causes cell cycle arrest and promotes the apoptosis of human neuroblastoma cells and in human neuroblastoma RRM2 is suppressed following chemotherapy. Oncology reports 40, no. 1: 355–360, 2018.

39. Kusari, J., U. S. Verma, J. B. Buse, R. R. Henry, and J. M. Olefsky. Analysis of the gene sequences of the insulin receptor and the insulin-sensitive glucose transporter (GLUT-4) in patients with common-type non-insulin-dependent diabetes mellitus. The Journal of clinical investigation 88, no. 4: 1323–1330, 1991.

40. Nisar Wani and Khliad Raza. Raw Sequence to Target Gene Prediction: An Integrated Inference Pipeline for ChIP-Seq and RNA-Seq Datasets. In: Malik H., Srivastava S., Sood Y., Ahmad A. (eds) Applications of Artificial Intelligence Techniques in Engineering. Advances in Intelligent Systems and Computing, 697, 2019. https://doi.org/10.1007/978-981-13-1822-1_52

